# Does the evolution of predatory behaviour alter stress response? Insights from a selection experiment on bank voles

**DOI:** 10.1101/2025.03.24.644903

**Authors:** Gokul Bhaskaran, Jella Wauters, Pawel Koteja, Edyta T. Sadowska

## Abstract

The ability to cope with challenging situations, such as the predator-prey interactions, can determine Darwinian fitness. While many studies concerned the stress response of prey facing a predator, the stress response of the predator remains understudied. We hypothesised that evolution of predatory lifestyle involves adaptive changes in the HPA axis regulation and tested this using an experimental evolution model comprising lines of bank voles (*Clethrionomys = Myodes glareolus*) selected for predatory behaviour towards crickets (P-lines) and unselected control lines (C-lines). We measured plasma corticosterone after a cricket hunting test and a sham test, where the voles experienced identical conditions but without prey. The test conditions affected corticosterone levels differently depending on the selection linetype and sex (significant interactions). After the sham hunting test, corticosterone level was significantly higher in P-lines than in C-lines, suggesting an increased alertness and mobilisation. Following the cricket hunting test, corticosterone levels tended to decrease in P-lines, but to increase in C-lines, although not significantly. These results support the hypothesis that selection for predatory behaviour has adaptively altered the HPA axis regulation. However, these effects were primarily driven by differences in females, highlighting the need to consider sex differences in studies of stress physiology.

## 1. Introduction

In nature, animals face a variety of stressful challenges such as food deficiency, extreme weather conditions, competitors or habitat disturbance, which require them to remain alert and adapt to these challenges [1]. One such situation is the predator-prey interaction. While many studies concerned the stress response of prey facing a predator [2–4], much less is known about the stress response on the predator side. Here we asked how the level of corticosterone, a component of the HPA axis stress response, changes in response to prey encounter, and whether and how the stress response has evolved in lines of a non-laboratory rodent, the bank vole (*Clethrionomys = Myodes glareolus*), selected for increased predatory behaviour towards crickets.

Predatory behaviour is important both ecologically and evolutionarily. It is a key driver of food webs, shapes species richness, and influences population dynamics [5–7]. Predation — defined as one organism killing another for food — has evolved independently several times (including several lineages of rodents [8,9]), often becoming a powerful evolutionary force [10]. Hunting is a challenging process involving movement in open habitats, where the risk of injury and exposure to other predators must be balanced against the energy costs and potential profits [11]. Successful coping with such challenges involves physiological processes that are activated or modulated as part of a stress response, which is typically accompanied by significant changes in stress hormone levels [12].

In vertebrates, the hypothalamic-pituitary-adrenal (HPA) axis acts as a key facilitator of stress responses and is vital in regulating homeostasis [13–15]. The HPA axis, through the secretion of glucocorticoids such as corticosterone, influences a broad range of an organism’s functions including metabolism, energy mobilisation, behaviour [16–19], and coordinates the adaptive response to stressors [20]. In the context of predatory behaviour, the role of HPA axis function warrants closer examination. Elevated glucocorticoids may have dual effects: they can enhance hunting activity by increasing energy availability and alertness, or they can inhibit performance by inducing distress.

Under natural conditions, numerous factors influence HPA axis activity, making it difficult to isolate changes due to specific selection pressures. However, experimental evolution provides a robust method for examining responses to selection for specific traits while controlling for random variation [21]. For example, golden hamsters (*Mesocricetus auratus*) were selectively bred for predatory behaviour based on locust capturing ability [22]. However, although several associated traits were analysed [23–25], the stress response of the hamsters was not studied. Mice, in which predatory behaviour towards crickets increased as a correlated response to selection for high voluntary wheel-running activity [26], have an increased baseline levels of corticosterone, suggesting the selection resulted in a modulation of HPA axis function [27,28], but analyses of corticosterone levels were not performed in the context of the predatory behaviour. Although stress-response functions have been analysed in several other selection experiments, either as directly selected traits [29,30] or as correlated responses to selection for other traits [31,32], none have directly addressed stress-response mechanisms associated with hunting.

We used a unique rodent model system: lines of bank voles selected for their ability to quickly catch crickets in a 10 min tests performed after a few hours of fasting (P – Predatory lines), and unselected Control (C) lines [33]. The effects of selection were visible from the third generation onwards. In generation 31, 62% of the P-line voles had caught the cricket already on the first encounter, and 84% had caught the cricket on at least one of the four trials, whereas only 5% of the C-line voles had caught the cricket on at least one of the replicated trails (Figure S1). Compared to C-line voles, the P-line ones exhibit proactive personality traits, such as increased activity and boldness, and moving faster on straighter trajectories [34], while spatial learning capabilities remained unaffected [35]. Lipowska *et al*. [36] reported that selection for the predatory behaviour did not affect baseline or post-acute restraint corticosterone levels, but the maximal response to ACTH stimulation was decreased, which implies a narrowed reaction scope and may mean that the relative response to the restraint stress was increased. In the P lines, successful hunting requires maintaining alertness despite a few hours fast in an empty cage prior to the test, and a proactive behaviour towards a novel, mobile object - the cricket - which, as we noticed, can trigger stress-related and evasion behaviour in some voles [36,37]. The selection experiment is therefore a good system to study how the stress response system adapts in the course of the evolution of predatory behaviour.

We hypothesised that the evolution of increased predatory propensity in P-line voles leads to altered HPA axis reactivity, resulting in differences in corticosterone levels during hunting compared to C-line voles. We measured glucocorticoids (target molecule: corticosterone) concentration in the vole’s plasma after a cricket hunting test such as applied in the selection protocol and after a sham hunting test, in which the animals faced the same stressful environment in the pre-test period (open space of a new, empty cage, and fasting), and without the cricket offered to hunt in the actual 10-min test. The corticosterone concentration measured after the sham hunting test can be regarded as a proxy for the corticosterone level at the start of the actual cricket hunting test. Because of the dual role and complexity of corticosterone [38], any specific predictions must be conditional.

We considered two hypothetical scenarios. 1) If an increased level of corticosterone reflects a distress, we expect that 1a) after the sham hunting test its level should be lower in P than in C lines (reflecting adaptation in P lines to the stressful environment), and 1b) after the cricket hunting test the level of corticosterone should further increase in C lines (cricket perceived as additional stressor) and not change in P-line voles (who should not perceive the cricket as a disturbing stressor). On the other hand, 2) if an increased level of corticosterone is a part of the mechanism responsible for increased alertness and mobilisation under the stressful conditions, we expect that 2a) after the sham hunting the corticosterone level will be higher in P than in C lines, and that 2b) after the cricket hunting test the level should decrease in P lines (a decreased “predatory drive” after the usually successful hunting) and not change in C lines (these voles usually do not respond actively to the crickets). The results of this experiment favoured the second scenario, but also showed that a predator’s HPA responses to hunting challenges can vary markedly with sex.

## 2. Materials and methods

### (a) Animals and the selection experiment

This work was performed on bank voles (*Clethrionomys = Myodes glareolus* Schreber 1780) from generation 35 of an ongoing artificial selection experiment [33,39]. The rationale, history and protocols of the selection experiment has been described comprehensively elsewhere [33,36,39]. Briefly, the colony was established with about 320 wild voles caught from Niepołomice Forest in southern Poland in 2000 and 2001. After 5-6 generations of random breeding, four replicate lines selected for predatory propensity (Predatory - P) and four randomly bred unselected Control (C) lines were created, with 15-20 reproducing families per line.

Animals were maintained in standard plastic mouse cages (mostly opaque, polypropylene) with sawdust bedding, at a constant temperature (20 ± 1°C) and photoperiod (16 h : 8 h light : dark; light phase starting at 2:00 am). Breeding pairs and pairs with offspring (up to 17 days old) were maintained in model 1290D cages (Tecniplast, Bugugiatte, Italy; dimensions L×W×H: 425×266×155 mm, floor area 800 cm^2^), equipped with a shelter, additional nest material (paper towels) and cardboard tubes (environment enrichment). At the age of 17 days the animals were weaned, marked temporarily by fur clipping and kept in family groups until the age of 30-35 days. At the age of about 34 days, all individuals were marked permanently with mouse ear tags (model 10005-1; National Band and Tag, Newport, KY; mass 0.18 g) and later maintained in same-sex groups of three individuals in model 1264C cages (dimensions L×W×H: 267×207×140 mm; floor area 370 cm^2^). Depending on the cleanliness, the cages were changed about every week. Water and food (a “breeding type” rodent chow: 24% protein, 3% fat, 4% fibre; Labofeed H, Kcynia, Poland) was provided *ad libitum*.

The predatory propensity was assessed in a cricket hunting test performed on adult animals (75-105 days of age), fasted before the trials. After a few hours of fasting a live cricket (*Gryllus assimilis*) was introduced to the cage and its presence and state were checked after 0.5, 1, 3, 6 and 10 min. The results were scored as ranks (rank 1–5: cricket caught in 0.5, 1, 3, 6, or 10 min, respectively; rank 6: cricket not caught). In the initial generations the voles were fasted for 10-12 hours before the tests, and the crickets were instars of about 1 cm length. In subsequent generations the fasting duration was gradually decreased (eventually to 2 hours), and the size of the crickets increased to about 2 cm. In the first two generations, three independent tests were performed (each on a different day). Later, the tests were performed on two days (with 7-10 days intervals), but two tests were performed on each day: one during light phase (about 15:00-16:00 hours) and the next after the beginning of dark phase of the colony photoperiod (18:00-19:00 hours), but with lights on. This eventual changes in selection protocol are detailed in Sadowska *et al*. [39]. The selection criterion was the mean of the ranked time to catch the cricket in the repeated tests, and ANCOVA-adjusted for sex, body mass and other cofactors.

### (b) Experimental design

A total of 193 adult voles were used in this experiment (96 males and 97 females, 12-13 males and females per each replicate C and P line). The animals were randomly sampled from the first and second litters of 12-14 families within each of the eight replicate lines (1-2 individuals per family). The animals had no contact with crickets. When the voles were at 80-90 days old (average age: 84.5), they were randomly assigned to two groups, in which either the standard cricket hunting tests (P lines: 48; C lines: 49) or a sham hunting tests (P and C lines: 48 each) were performed. Animals from the same family were not assigned to the same test type. The tests were performed in blocks of about 8 individuals per day with nearly equal representation of sex, selection direction and test type.

### (c) Hunting tests, video recordings and behavioural analysis

The hunting tests were performed in a similar way as the test used for selection protocol. The tests were conducted during day time (between 11:30-15:00 hours CET). The voles were picked from their native maintenance cage, weighed (± 0.01 g) and transferred to a fresh cage of the same type as used in regular maintenance. No fresh bedding was provided, and only a small amount of sawdust from its native cage was moved to the new one (to maintain olfactory cues). No food was provided, but water was available *ad libtum*. After 2 hours, the cage lid and water bottle were replaced with a tight, vertically oriented open box made of thick white paper sheets, extending the cage walls to 22 cm, which prevented the voles from escaping while allowing undisturbed camera recordings. This cage was then moved to the testing room and either of the 10-min hunting test began immediately. At the beginning of the cricket hunting test, a live cricket (adult, about 2 cm long) was placed in the cage. In sham hunting test, no cricket was added, instead just only the hand gesture was made such as when placing the cricket.

The tests were video recorded (resolution 704 × 576 pixels, 25 frames/sec) for 10 min using a wired camera (SAMSUNG SCB-3000P) mounted centrally 100 cm above the cage bottom, connected to digital recorder (BCS, Warsaw, Poland). The recording system was operated from outside the testing room and a monitor screen facilitated real-time supervision during the test.

Each test was marked with a unique ID and recordings were analysed by an observer blind to the vole’s selection linetype. We measured two parameters: the cricket capture status and the time to catch the cricket in a successful test (time measured in seconds). From the video recordings of the cricket hunting test, a trial was classified as a “successful” if the vole caught and killed the cricket, or as “not successful” if the cricket remained alive at the end of the test.

### (d) Blood collection, sample processing and corticosterone level analysis

Immediately after the test, the vole was removed from the cage and approximately 75 µl of blood was collected from the retro-orbital venous sinus using sterile heparinized capillary tubes (BR749321, MERCK). Retro-orbital sampling was chosen as the only method that could guarantee sufficient volume of blood from live bank voles. Blood sampling took no more than 3 min (mean ± standard deviation: 1:22 ± 0:29 min), which presumably has negligible effect on corticosterone levels [40], later confirmed by statistical analysis. Anaesthesia was not applied prior to blood sampling, as it could impair the ability to obtain samples of sufficient volume through reduction of blood flow. Moreover, inducing anaesthesia would prolong the blood sampling procedure, which would compromise measurement of corticosterone levels [41]. The blood sampling was performed outside the test room. The capillary tubes containing blood samples were transferred to Eppendorf tubes (1.5 ml) and stored on ice until samples were collected from all animals in a block.

Plasma was separated by centrifugation (15 min at 14000 g), transferred to fresh Eppendorf tubes, and stored at −80°C. Later, these plasma samples were transported to the Leibniz Institute for Zoo and Wildlife Research (Berlin) for analysis of glucocorticoids (target molecule: corticosterone). Because of cross-reactivities of the antibody we could not exclude the possibility that the reported concentration includes also some corticosterone metabolites, but, unlike in the case of the non-invasive measurements performed on faeces, this bias is presumably very small [42].

Depending on the available plasma volume, either 90 µl or 95 µl of distilled water was added to 10 µl or 5 µl of plasma, respectively. Extraction was then performed using 2 ml of a tert-methylbutylether/petroleum ether (TBME/PE) mixture (30:70 v/v) by shaking for 30 min. After freezing at 70°C for 15 min the organic phases were decanted, dried down at 55°C for 10 min under nitrogen and reconstituted in 0.25 ml of 40% methanol. Corticosterone was measured as described by Dehnhard *et al*. [43], using a polyclonal antibody (rabbit) against 4-pregnen-11β,21-diol-3,20-dione-21-BSA, corticosterone-peroxidase as label and corticosterone as standard. The antibody was verified and showed the following cross-reactivities: 4-pregnen-11β,21-diol-3,20-dione (corticosterone), 100%; 4-pregnen-21-ol-3,20-dione (deoxycorticosterone), 24.4%; 4-pregnen-11α,17,21-triol-3,20-dione (hydrocortisone), 13.4%, and 4-pregnen-3,20-dione (progesterone), 21.8%.

The sensitivity of assays was defined as two standard deviations from the signal given by the zero blank and was 0.2 pg/well, which allows detection with a limit of 3.75 ng ml^−1^ [43]. The range of the calibration curve was 0.8-400 pg/20µl. Serial dilutions of plasma extracts yielded displacement curves parallel to standard corticosterone. The intra-assay CV, determined on two biological samples including low and high concentration respectively (16 repeats in duplicate each), was 17.8% and 8.5%, respectively. The linear range, normally between B80 and B20, was therefore adapted to 20-130 pg/20 µl.

The inter-assay CV (12 assays), based on a low-quality control sample (LQC) and high-quality control sample (HQC), both fitting the (adapted) linear range of the curve and run in duplicate, was respectively 6.0 and 7.3%. All EIA measurements were performed in duplicate with acceptance criteria of a coefficient of variation (CV) below 5%.

### (e) Statistical analyses

To assess the difference in the proportion of voles capturing the cricket between the P and C lines, we used logistic regression model with selection direction and sex as fixed factors and the capture status as binary response variable. Replicate line nested within selection direction was included as a random effect. The analysis was performed with SAS v. 9.4 Glimmix procedure (SAS Institute Inc., Cary, NC, USA), with RSPL method of estimation and variance components restricted to non-negative values.

Further analyses were performed with SAS Mixed procedure, with REML method of estimation and variance components restricted to non-negative values. Body mass and corticosterone concentration was tested using cross-nested ANCOVA models with three main fixed factors: test type (sham and cricket hunting), selection direction, and sex. Replicate line nested within selection direction was included as a random effect. The hierarchical structure of the statistical model (replicate lines nested in selection direction) is required to allow a proper distinction of the effects of selection from random genetic effects, such as genetic drift [44]. The selection direction × test type and selection direction × sex were always included as they were biologically relevant and meaningful for testing the hypothesis, but other interactions between main factors were removed if they were not significant (p > 0.05). If interaction between fixed categorical factors was significant, the effect of one of the fixed factors was tested separately for each level of the other (but within the framework of the same model, using “slice” option of SAS Mixed procedure). Additionally, we included litter number as a cofactor, and age, test date and test time, as covariates. For the corticosterone model, body mass and time to sample blood were included as additional covariates. The initial model for the corticosterone concentration included all main factors interaction with body mass, but these were removed as non-significant. If the absolute value of studentized residual was higher than 3.5, it was considered as an outlier. To achieve normality of the residual distribution, the corticosterone concentration was log10-transformed and two outliers were removed (both were from cricket hunting test, one C-line: residual value = -4.17, and one P-line: residual value = -3.57).

As only two C-line voles captured the cricket, the analysis of the time to catch the cricket and its partial correlation with corticosterone concentration was restricted to voles from P lines that caught the cricket. For the analysis of time to catch the cricket as dependent variable, we used mixed-ANCOVA models with sex as the fixed factor and replicate line as a random effect. Litter number was included as a cofactor. In the initial model, we included age, body mass, test date and test time as covariates. A similar model was used for the analysis of partial correlation between corticosterone concentration (the dependent variable) and the time to catch the cricket (one of predictors). Time of sampling blood was included as an additional covariate. In both analyses, the covariates were sequentially removed if their effects were not significant. To achieve normality of the residual distribution, both corticosterone concentration and time to catch a cricket were log10-transformed.

Complete tables with results from the models are presented in Supplementary materials. Here, we provide the main results as adjusted least-square means with 95% confidence interval (LSM ± CI).

## 3. Results

Body mass ranged from 14.31 to 35.29 g (mean ± s.d.: 22.16 ± 4.02 g). Males were heavier than females (*p* < 0. 0001, Figure 1a, Table S1). Body mass did not differ between the Predatory (P) and Control (C) lines (*F*_1,6_ = 0.11, *p* = 0.75) or between the voles assigned to cricket or sham hunting test (*F*_1,176_ = 0.47, *p* = 0.49). The age, test date, time of day, and litter number did not influence body mass (*p* ≥ 0.51).

**Figure 1.**
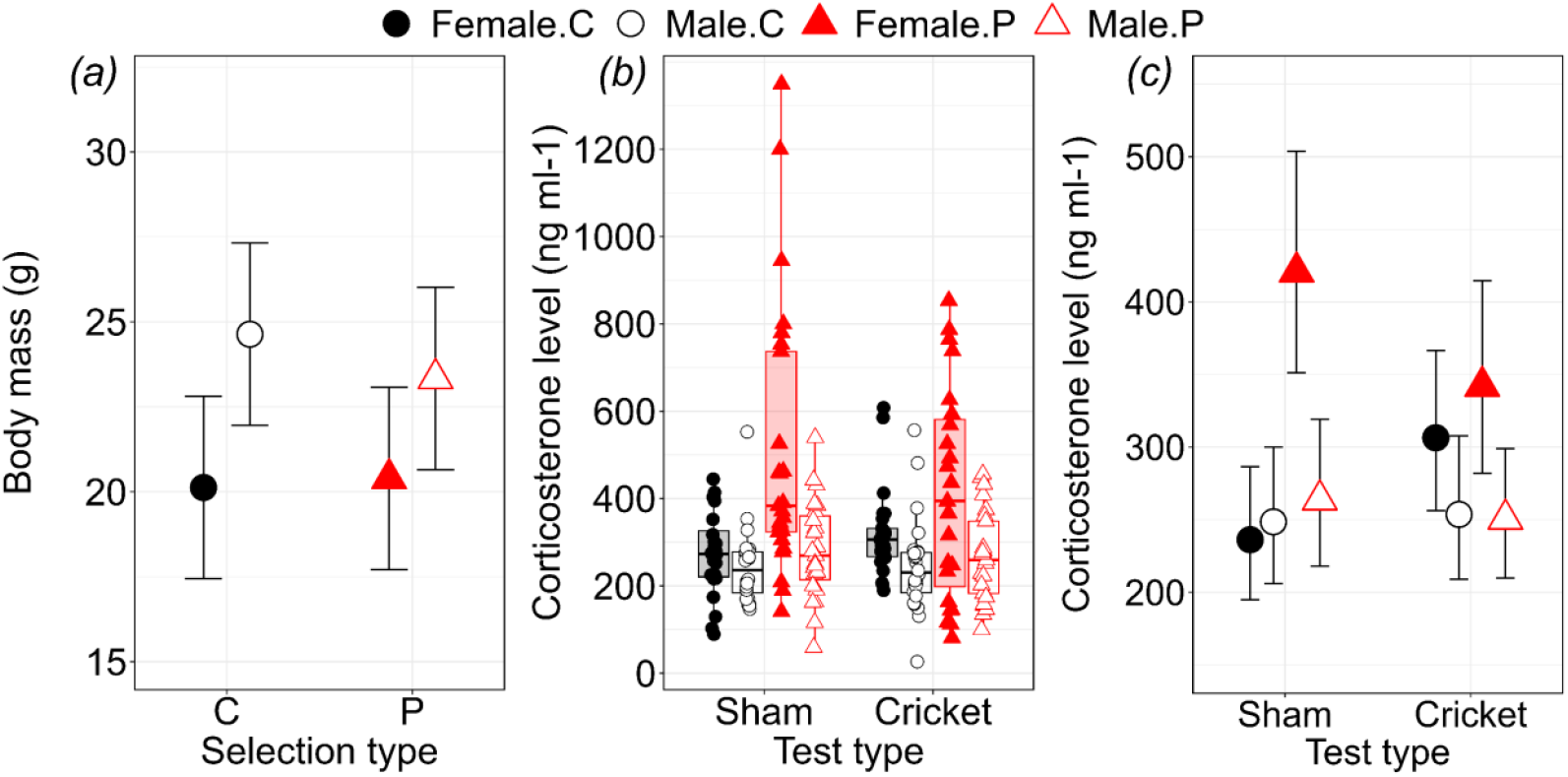
a) Body mass (g) of female and male bank voles from Control and Predatory lines (adjusted least-square means from mixed ANCOVA models ±95% Cls). b) Corticosterone concentration (ng ml^−1^) of female and male bank voles from Control and Predatory lines after sham and cricket hunting tests. Boxes represent Inter Quartile Range (IQR); median lines are at Q2; whiskers show the range from Q1-1.5×ICR to Q3 +1.5×IQR; and the points represents raw values. b) Corticosterone concentration (ng ml^−1^) of female and male bank voles from Control and Predatory lines after sham and cricket hunting tests (Adjusted least-square means from mixed ANCOVA models ±95% CIs. The values for the plot were taken from the model that included three-way interaction: test type × selection direction × sex (the values reported in the text, which were the basis for statistical inferences, are from the reduced model without this three-way interaction). The values are back-transformed from the analyses on log10-transformed data. Open symbols denote males, and filled symbols females.

In the cricket hunting test, 83% of the P-line voles caught the cricket (40 out of 48; 5 males and 3 females did not catch the cricket), whereas only 4% of the C-line voles caught the cricket (2 out of 49; 1 male and 1 female; effect of selection in logistics regression model: *F*_1,94_ = 32, *p* < 0.0001). In P lines, the time to catch the cricket did not differ between males and females (linear mixed model: *F*_1,35_ = 0.32, *p* = 0.57; Figure S3). Voles from the first litter captured the crickets faster those from the second litter (*F*_1,34_ = 4.8, *p* = 0.035). Age, body mass, the test date and time of day did not significantly affect the time to catch the cricket (*p* ≥ 0.14).

The corticosterone concentration ranged from about 20 to 1350 ng ml^−1^ (Figure 1b, Figure S2) and decreased with body mass of individuals (slope ± s.e.: -0.0092 ± 0.0041; *F*_1,179_ = 5.1, *p* = 0.025). The mass-adjusted concentration did not differ systematically between that measured after the cricket and the sham hunting tests (*F*_1,179_ = 0.01, *p* = 0.92; Figure 1c, Table S1). The corticosterone level was on average higher in P than in C lines (*F*_1,179_ = 7.9, *p* = 0.005, Figure 1c, Table S1) and higher in females than in males (*F*_1,179_ = 9.3, *p* = 0.003), but the results were complicated by significant interactions of the effect of selection with the hunting test type (*F*_1,179_ = 4.2, *p* = 0.04) and with sex (*F*_1,179_ = 6.0, *p* = 0.015; Figure 1c, Table S1). After the sham hunting test, the corticosterone concentration was on average higher in P than in C lines (*F*_1,179_ = 12, *p* = 0.0007). Compared to these values, the corticosterone level after the cricket hunting test tended to decrease in P lines but to increase in C lines (although these effects were not significant: p > 0.4), so that after the cricket hunting test there was no difference between the P and C lines (*F*_1,179_ = 0.33, *p* = 0.56; Figure 1c). However, as already mentioned, the results were also affected by the interaction between the selection direction and sex (*F*_1,179_ = 6.0, *p* = 0.015). In females, the corticosterone concentration was higher in P lines than in C lines (*F*_1,179_ = 14, *p* = 0.0002), whereas in males there was no effect of selection (*F*_1,179_ = 0.09, *p* = 0.77). The corticosterone concentration was higher in females than in males in P lines (*F*_1,179_ = 16, *p* < 0.0001), but not in C lines (*F*_1,179_ = 0.48, *p* = 0.49). Litter number, age, the test date, and time of day did not significantly affect the corticosterone concentration (*p* ≥ 0.27). Importantly, the time to sample blood did not significantly affect the corticosterone concentration (*F*_1,179_ = 0.03, *p* = 0.86). In P-line voles that successfully captured the cricket, the corticosterone concentration was not correlated with the time to catch the cricket (*F*_1,34_ = 0.89, Figure S3, *p* = 0.35).

## 4. Discussion

As expected, voles from the lines selected for predatory behaviour (Predatory, P lines) captured crickets more often and faster than those from the unselected Control (C) lines. The corticosterone concentration ranged from about 20 to 1350 ng ml^-1^ (Figure 1b) and was independent of body mass, but was differentially affected by the test conditions depending on the selection linetype and sex of the voles (significant interactions; Figure 1c). After the sham hunting test, i.e. under conditions similar to those experienced by voles at the start of the cricket hunting test used in the selection protocol (after a few hours in an open space of a new, empty cage without food), the corticosterone level was significantly higher in the P lines than in C lines. Compared to this level, the corticosterone concentration after the cricket hunting test tended to decrease in the P lines, but to increase in the C lines (although these tendencies were not significant and appeared only in females). Consequently, the corticosterone concentration after the cricket hunting test did not differ significantly between the P and C lines.

We considered two hypothetical scenarios of adaptive changes in the HPA regulation, related to the dual role of corticosterone. First, if the level of corticosterone is primarily a signal of how the animal is ‘stressed’ (and a high level would indicate ‘distress’), the corticosterone concentrations after both the sham and cricket hunting tests should be lower in voles from the P lines than those from C lines, because the selection should have resulted in better coping with the stressful situation (being placed in a new, empty cage without food). The results of our experiment clearly rejected this scenario, as corticosterone levels were higher in the P lines. Second, if the level of corticosterone is primarily a signal of the animal’s increased metabolic mobilisation, alertness and readiness to proactively cope with the stressful conditions, for the same reason the corticosterone concentrations after the tests should be higher in voles from the P lines. Additionally, if the level of corticosterone is particularly associated with the predatory propensity, we expected that the corticosterone concentration in P lines should be lower after cricket hunting test than after the sham hunting test, as the successful hunting should decrease the “predatory drive”. The results provide a convincing support for the first part of this hypothesis, but only a weak for the second part.

A question arises, which specific physiological, biochemical and neurobiological mechanisms underly such adaptive (in the context of the experimental evolution model) behavioural changes in the P lines. Voles from P lines not only show predatory behaviour in the laboratory cricket hunting test, but also prefer animal-based food under semi-natural conditions, as evidenced by their higher hair δ^15^N values than C lines [45], indicating that selection has led to the evolution of a predatory lifestyle. They also show several other characteristics of active hunting predators, including higher food consumption and increased home cage activity [46]. P-line voles tend to have a higher basal metabolic rate (BMR) compared to C-line voles (although not statistically significant; [39]), consistent with comparative analyses across mammals showing that predators have on average higher BMR [47]. In the open field test, P-line voles moved faster and covered more distance on relatively straighter trajectories compared to those from the C lines [34]. Transcriptomic analyses showed that the selection resulted in changes in allele frequencies or expression levels of genes associated with the regulation of hunger and aggressive behaviour, as well as cAMP activity and GABAergic, dopaminergic and serotonergic transmission, which can in turn affect locomotor activity, arousal and attention [48]. Thus, the evolution of predatory lifestyle in P-line voles involves two key aspects: a) higher energy demand and b) a more proactive personality to support active hunting. These two factors could explain the corticosterone differences between the sham and cricket hunting tests.

Due to their higher energy requirements, P-line voles presumably need to feed more frequently than the C-line voles. Consequently, after the same fasting period before the hunting tests, they are likely to exhibit a greater appetite compared to C-line voles. This heightened drive for food acquisition is consistent with their proactive personality, which includes increased locomotion, exploration, and a greater tendency to engage in persistent goal-directed activities. Such traits are commonly associated with a more active foraging strategy, where energy must be efficiently mobilised to support movement and hunting efforts [49]. Corticosterone plays a crucial role in this process by facilitating metabolic mobilisation through gluconeogenesis, lipolysis, and proteolysis, ensuring a continuous supply of energy during periods of increased activity [50–53]. Thus, the upregulation of the HPA axis could lead to increased corticosterone release, which in turn may enhance the animal’s activity levels. One potential mediator of such a relationship is corticotropin-releasing factor (CRF). CRF stimulates corticotropin and glucocorticoid secretion, but also has behavioural effects in the central nervous system [54], independent of its action on the anterior pituitary. It is also possible that corticosterone itself plays a role in promoting active movements, as has been reported in other vertebrates [55,56]. Corticosteroids play an important role in fear and anxiety, enabling behavioural adaptation by facilitating information processing in potentially dangerous or stressful environments (for a review, see [57]). We are also aware of studies that have reported a reduction in activity levels following experimental elevation of corticosterone levels [58–60]. However, our results imply that the elevated corticosterone level in P-line voles reflects HPA axis activity linked to predatory lifestyle, which underlies heightened alertness and readiness to proactively cope in a stressful situation.

The observed effects were primarily driven by difference observed in females, while corticosterone levels in males did not appear to be substantially affected by selection or test conditions. Thus, the effect of selection on HPA axis activity appears to be stronger in females than in males. This suggests that physiological responses to predation-related contexts may be sex-dependent, emphasizing the importance of considering sex as a critical factor in studies of stress physiology and behavioural adaptation. As the selection experiment on bank voles continues, the specific characteristics of the HPA axis activity and signalling can be addressed in further studies.

## 5. Conclusions

Our study has shown that the experimental evolution of an active predatory lifestyle in a rodent, the bank vole, is associated with increased corticosterone levels, which enhance metabolic mobilisation and alertness under the challenging conditions encountered during hunting. Future research on this model should continue to explore the interaction between behaviour and physiological responses, considering both the broader ecological context and sex differences, as well as the lower-level molecular and neurophysiological mechanisms on the other.

## Supporting information

A separate file that includes supplementary materials (three figures and one table)

## Ethics

The animal colony was under the supervision of a qualified veterinary surgeon. All the experimental procedures were approved by the decisions of the Local Bioethical Committee in Cracow, Poland (selection experiment: No. 258/2017; hunting tests and blood sampling: No. 186/2022).

## Data accessibility

The dataset and code associated with this manuscript can be found at [61].

A separate file that includes supplementary materials (three figures and one table).

## Declaration of AI use

We have not used AI-assisted technologies in creating this article.

## Authors’ contributions

G.B: Data curation, formal analysis, investigation, methodology, validation, visualization, writing-original draft, writing-review & editing; J.W: Investigation, validation, writing-review & editing; P.K: Conceptualization, formal analysis, funding acquisition, methodology, resources, software, supervision, validation, writing-review & editing; E.T.S: Conceptualization, formal analysis, funding acquisition, methodology, project administration, resources, software, supervision, validation, writing-review & editing.

All authors gave final approval for publication and agreed to be held accountable for the work performed therein.

## Conflict of interest declaration

We declare we have no competing interests.

## Funding

The project was supported by the National Science Centre 2020/39/B/NZ8/02996 (to ETS). The project was also supported by the Jagiellonian University funds: N18/DBS/000003 and DS/WB/INOS/757. The funding source did not affect the experimental design, data analysis or writing the manuscript.

## Acknowledgments

The authors would like to thank Katarzyna Baliga-Klimczyk, who managed the animal colony. We are grateful to Natalia Boron for her help with the video analyses. We also want to thank Małgorzata M. Lipowska and Alaa Hseiky for their comments and suggestions on an earlier version of the manuscript.

## Notes

https://doi.org/10.57903/UJ/LBPCGT

### Competing Interest Statement

The authors have declared no competing interest.

