## Supplementary material for "Does the evolution of predatory behaviour alter stress response? Insights from a selection experiment on bank voles": A separate file that includes supplementary materials (three figures and one table)

### Supplementary figures

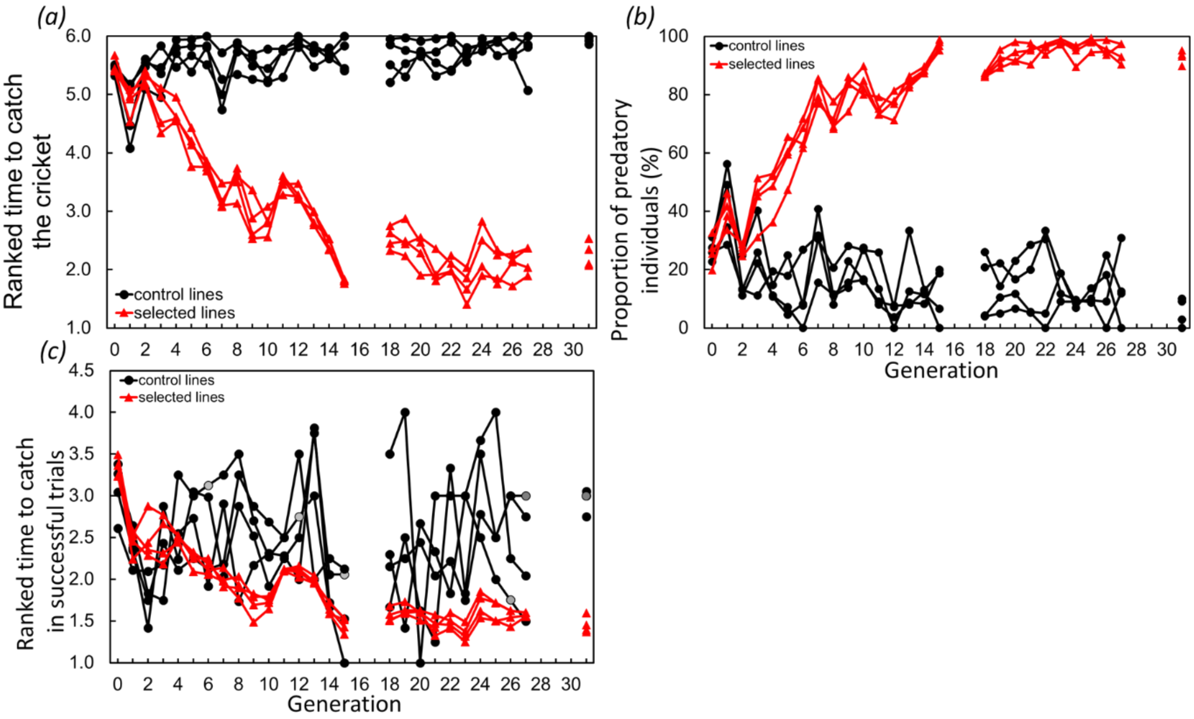


Figure S1: Direct phenotypic responses to 31 generations of selecting bank voles towards (a) predatory propensity measured as ranked time to catch a cricket (the selection criterion), (b) proportion of predatory individuals (in percentage) and (c) ranked time to catch cricket in successful trials (grey points for C lines indicate cases where no individual caught the cricket; these values are estimates based on the mean of the values from the previous and next generation or the values from the previous generation if subsequent data are unavailable). The gaps in between generations were relaxed selection.

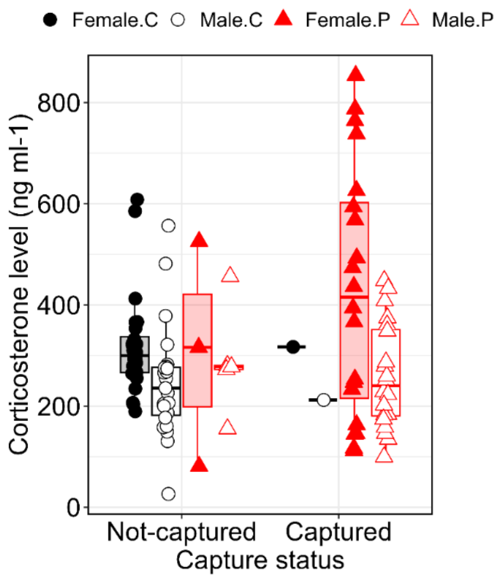


Figure S2. Corticosterone concentration of female and male bank voles from Control and Predatory lines that not-captured and captured crickets. Box represents IQR (Inter Quartile Range); median line at Q2; whiskers show Q1/Q3 ±1.5×IQR and the points represents raw value.

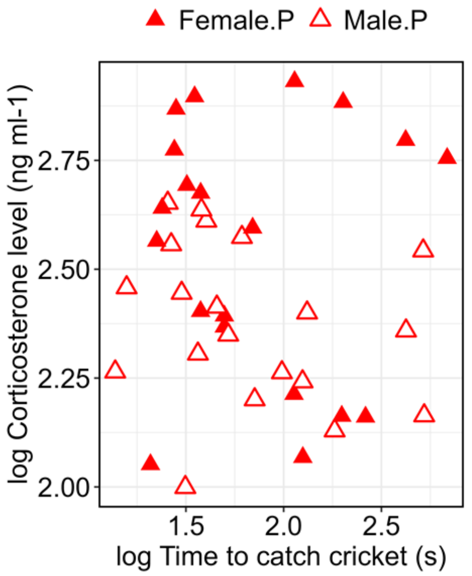


Figure S3. Corticosterone concentration of female and male bank voles from Predatory lines measured after cricket hunting test in relation to the time to catch the cricket (both variables are log10-transformed) after cricket hunting tests. Open symbols denote males, and filled symbols females.

### Supplementary table

Table S1. Results from from the Linear Mixed ANCOVA models: tests statistics *F* (d.f.) = F-value (denominator df), *P* = significance levels, LSM ± CI = Least Square Means ± Confidence Interval. The values reported for corticosterone and time to catch cricket traits are from analyses after log-10 transformation.

| Factors in model | Levels | | Traits | | |
|---|---|---|---|---|---|
| | | | Body mass (g) | Corticosterone (ng/ml) | Time to catch cricket (s) |
| test type | | *F* (d.f.) | 0.47 (176) | 0.01 (179) | |
| | | *P* | 0.49 | 0.92 | |
| | cricket hunting | LSM ± CI | 22.3 ± 1.9 | 2.46 ± 0.04 | |
| | sham hunting | LSM ± CI | 22.0 ± 1.9 | 2.45 ± 0.04 | |
| selection direction | | *F* (d.f.) | 0.11 (6) | 7.9 (179) | |
| | | *P* | 0.75 | 0.005 | |
| | P | LSM ± CI | 21.9 ± 2.7 | 2.50 ± 0.04 | |
| | C | LSM ± CI | 22.4 ± 2.7 | 2.41 ± 0.04 | |
| sex | | *F* (d.f.) | 73 (176) | 9.3 (179) | 0.32 (35) |
| | | *P* | <0.001 | 0.003 | 0.57 |
| | female | LSM ± CI | 20.3 ± 1.9 | 2.51 ± 0.04 | 1.90 ± 0.34 |
| | male | LSM ± CI | 24.0 ± 1.9 | 2.40 ± 0.04 | 1.82 ± 0.34 |
| selection direction x test type | | *F* (d.f.) | 1.6 (176) | 4.2 (178) | |
| | | *P* | 0.21 | 0.042 | |
| | cricket hunting - P | LSM ± CI | 21.7 ± 2.7 | 2.47 ± 0.06 | |
| | sham hunting - P | LSM ± CI | 22.0 ± 2.7 | 2.52 ± 0.06 | |
| | cricket hunting - C | LSM ± CI | 22.8 ± 2.7 | 2.44 ± 0.06 | |
| | sham hunting - C | LSM ± CI | 22.0 ± 2.7 | 2.38 ± 0.06 | |
| selection direction x sex | | *F* (d.f.) | 3.3 (176) | 6.0 (179) | |
| | | *P* | 0.069 | 0.015 | |
| | female - P | LSM ± CI | 20.4 ± 2.7 | 2.58 ± 0.06 | |
| | male - P | LSM ± CI | 23.3 ± 2.7 | 2.41 ± 0.06 | |
| | female - C | LSM ± CI | 20.1 ± 2.7 | 2.43 ± 0.06 | |
| | male - C | LSM ± CI | 24.6 ± 2.7 | 2.40 ± 0.07 | |
| litter number | | *F* (d.f.) | 0.04 (177) | 1.2 (179) | 4.8 (34) |
| | | *P* | 0.85 | 0.27 | 0.035 |
| | first | LSM ± CI | 22.0 ± 1.9 | 2.48 ± 0.06 | 2.01 ± 0.34 |
| | second | LSM ± CI | 22.2 ± 1.9 | 2.43 ± 0.05 | 1.71 ± 0.34 |
| age (days) | | *F* (d.f.) | 0.44 (177) | 0.00 (179) | |
| | | *P* | 0.51 | 0.96 | |
| time of day | | *F* (d.f.) | 0.21 (177) | 0.01 (179) | |
| | | *P* | 0.65 | 0.93 | |
| test date | | *F* (d.f.) | 0.0002 (178) | 0.27 (179) | |
| | | *P* | 0.99 | 0.60 | |
| body mass (g) | | *F* (d.f.) | - | 5.1 (179) | 2.2 (36) |
| | | *P* | - | 0.025 | 0.15 |
| time to sample blood (s) | | *F* (d.f.) | - | 0.03 (179) | |
| | | *P* | - | 0.86 | |
